# DeviceAgent: An autonomous multimodal AI agent for flexible bioelectronics

**DOI:** 10.1101/2025.10.10.681748

**Authors:** Jaeyong Lee, Zuwan Lin, Wenbo Wang, Jongmin Baek, Ariel J. Lee, Almir Aljović, Arnau Marin-Llobet, Xinhe Zhang, Ren Liu, Na Li, Jia Liu

**Affiliations:** John A. Paulson School of Engineering and Applied Sciences, Harvard University, Boston, MA, USA; Broad Institute of MIT and Harvard, Cambridge, MA, USA; Electrical Engineering and Computer Science, MIT, Cambridge, MA, USA

**Author notes:** These authors contributed equally (Jaeyong Lee, Zuwan Lin, Wenbo Wang, Jongmin Baek).

## Abstract

The development of flexible bioelectronics remains a complex, multidisciplinary process that demands specialized expertise and labor-intensive efforts, limiting scalability, adaptability and accessibility. Here, we introduce DeviceAgent, an autonomous multimodal AI agent that integrates large language models (LLMs), vision-language models (VLMs), and domain-specific computational tools into a unified framework for bioelectronics research. Leveraging the emergent reasoning abilities of LLMs and VLMs, DeviceAgent enables zero and few-shot generalization, contextual learning, and flexible task execution across modalities. A multimodal context memory system orchestrates these capabilities, providing end-to-end support across the experimental pipeline–from high-level design objectives to fabrication protocol generation, visual defect inspection, and electrophysiological signal analysis, while maintaining human oversight at critical decision points. We demonstrate its capabilities through the development of stretchable mesh electronics for interfacing with human induced pluripotent stem cell-derived cardiomyocytes (hiPSC-CMs), a representative application involving complex device architectures, heterogeneous material nanofabrication, and electrophysiology analysis. DeviceAgent autonomously (1) generates customized bioelectronic layouts; (2) creates comprehensive fabrication protocols tailored to specific materials and processes; (3) identifies microscopic defects using visual reasoning; and (4) analyzes cardiac electrophysiological recordings in an interpretable manner. By embedding LLMs and VLMs within a structured, tool-augmented architecture, DeviceAgent establishes a scalable and accessible paradigm for AI-scientist collaboration in nanofabrication and bioelectronics research.

## Main

Recent advances in flexible and stretchable electronics have enabled transformative applications ranging from wearable health technologies to biomedical implants^1,2,3^. By combining miniaturized electrical components with mechanically compliant substrates, these systems conform to soft, dynamic biological systems, reducing mechanical mismatch and improving biocompatibility. Flexible bioelectronics enable chronic, high-resolution interfacing with biological systems, opening opportunities in fundamental biology, disease modeling, and therapeutic development^4,5,6,7,8^.

Despite this progress, the development of flexible bioelectronics remains a fragmented and labor-intensive process. A typical workflow requires translating biological or clinical goals into device concepts, designing multilayer layouts, generating computer-aided design (CAD) masks for layer-by-layer photolithography, tailoring fabrication pipelines to specific materials and architectures, and finally integrating devices with biological systems for testing. Each stage demands combination of domain-specific expertise in diverse fields like electrical engineering, materials science, or biology, and small errors can propagate across interdependent steps. Existing tools provide limited support: general-purpose CAD software lacks bioelectronics-specific features, fabrication protocols rely on ad hoc knowledge and manual optimization, and data analysis remains a separate, code-intensive process requiring expert interpretation. While electronic design automation (EDA) has transformed silicon circuits^9^, it offers limited support for hybrid materials and biological constraints, leaving researchers to bridge gaps between design, fabrication, and analysis with fragmented, manual processes.

Recent advances in large language models (LLMs)^10^ and vision-language models (VLMs)^11^ offer promising solutions to address these bottlenecks. These models can process and generate multimodal information including text and images, with emergent capabilities such as contextual reasoning, in-context learning, and zero-shot generalization^12,13,14^. Their adoption across scientific domains has already enabled autonomous workflows in diverse fields such as neural computation^15^, spatial biology^16^, behavior analysis^17^, and chemical synthesis^18^. In principle, LLMs could be applied to interpret fabrication recipes or generate code for device layouts, while VLMs could potentially detect microscopic defects or assess signal quality. Yet, LLMs/VLMs can only produce outputs in response to prompts. To execute end-to-end experimental workflows, models need to be embedded within an agent that provides orchestration, memory, and persistence, enabling it to define questions, act on datasets, and coordinate tasks across tools and environments.

To address this, we developed DeviceAgent, an autonomous multimodal AI agent that embeds LLMs and VLMs within a structured, tool-augmented framework. DeviceAgent connects reasoning models to specialized tools for CAD design, fabrication protocol generators, visual inspection, and signal analysis, all coordinated by an orchestration layer that maintains context, tracks progress, supports multi-step reasoning, and takes actions. Unlike conventional software that rely on isolated prompts and specific tasks, DeviceAgent provides end-to-end support across the design-fabrication-analysis pipeline while preserving human oversight at critical decision points.

We demonstrate DeviceAgent’s capabilities through a representative use case: the development of stretchable mesh electronics for interfacing with human-induced pluripotent stem cell-derived cardiomyocyte tissues (hiPSC-CMs)–a demanding application that involves precise device geometry, multi-material fabrication, and complex electrophysiological analysis^19,20^.

### DeviceAgent architecture and workflow

The development of bioelectronics has traditionally relied on a sequential, expertise-intensive workflow (**Fig. 1a**), progressing through discrete steps: defining objectives, designing device structures, generating CAD layouts, creating fabrication protocol, executing nanofabrication, and finally integrating with biological systems for electrophysiological analysis. Key tasks such as electrode placement, interconnects routing, and input/output (I/O) pad configuration are typically performed manually, requiring technical proficiency in both mechanical and electrical design principles along with a deep understanding of biological constraints. All the steps demand specialized knowledge across materials science, mechanical engineering, electrical engineering, or biology, making the process labor-intensive and inaccessible to newcomers.

**Fig 1.**
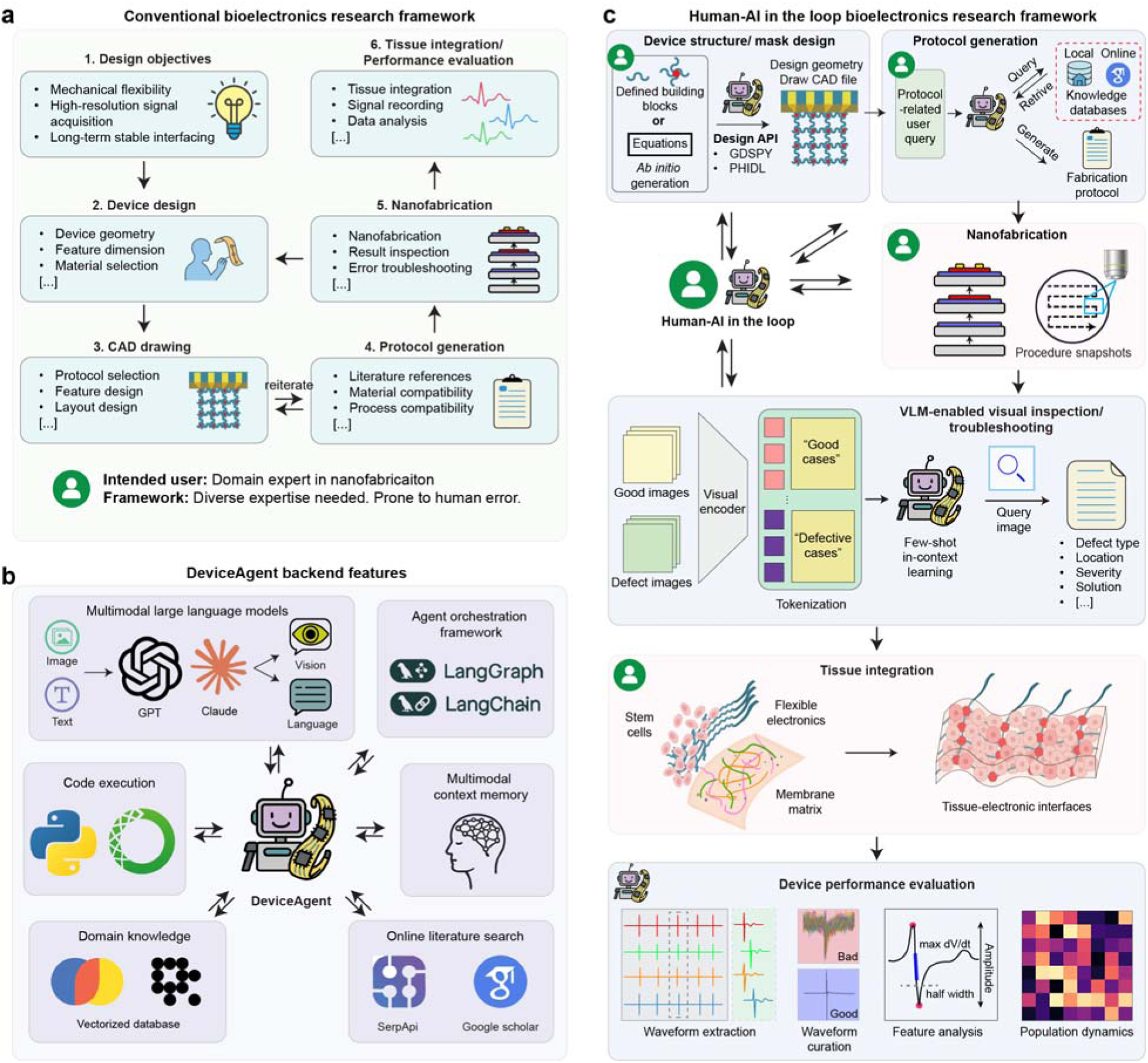
DeviceAgent system architecture. (**a**) The conventional bioelectronics research workflow follows a sequential process that requires specialized expertise at each stage, from defining objectives through device design, CAD drafting, and protocol generation to nanofabrication, tissue integration, and data analysis. This approach demands diverse expertise and is prone to human error. (**b**) The backend architecture of DeviceAgent integrates multimodal LLMs (GPT, Claude), agent orchestration frameworks (LangGraph, LangChain), code execution modules, multimodal context memory, a domain knowledge database, and online literature retrieval functions. (**c**) The human-AI-in-the-loop bioelectronics research framework proposed by DeviceAgent transforms each stage of the workflow: design geometry and mask creation using Python packages, protocol generation combining local and online knowledge, nanofabrication with VLM-enabled visual inspection, tissue integration, and comprehensive data analysis. This collaborative approach preserves human oversight while automating technical tasks.

Moreover, this fragmented workflow is susceptible to cumulative human error and often requires multiple design-fabrication-analysis cycles. Even simple designs require significant time investment, while complex layouts further elevate these demands. Fabrication protocol development introduces additional bottlenecks, often relying on extensive literature review and fabrication expertise to ensure material and process compatibility.

To address these limitations, we developed DeviceAgent, an autonomous multimodal AI agent designed to streamline bioelectronics design, fabrication, and evaluation (**Fig. 1b**). Its backend architecture integrates several key components: (1) multimodal LLMs, such as GPT-4o^21^ and Claude-3.7-Sonnet^22^, which function as reasoning engines capable of processing input, reasoning, defining customized workflows, and interacting with tools to accomplish them; (2) an agent orchestration framework empowered by LangGraph and LangChain^23^ that coordinates complex, multi-step reasoning; (3) a CodeAct-inspired code execution environment powered by Python that enables real-time implementation of computational tasks^24^; (4) a multimodal context memory system that maintains coherence across extended interactions; (5) a domain knowledge database containing vectorized information about bioelectronics device structures, materials properties, and fabrication techniques^25^; and (6) an online literature search capability using SerpAPI^26^ and Google Scholar that provides access to the latest research findings.

We designed the architecture to demonstrate a flexible, human-AI-in-the-loop pipeline (**Fig. 1c**) that enables flexible, iterative collaboration between researchers and the AI agent throughout the entire process. Specifically, during layout and mask generation, researchers can define high-level parameters in natural language descriptions or allow DeviceAgent to autonomously generate layouts using mathematic formulations. DeviceAgent translates these specifications into precise CAD designs using Python packages such as GDSPY^27^ and PHIDL^28^. For fabrication protocol generation, DeviceAgent combines internal knowledge databases with real-time literature retrieval to create comprehensive fabrication instructions. During experimental execution, when researchers use the agent-generated masks and protocols for fabrication, DeviceAgent provides visual inspection support through VLM-based defect detection, identifying defect type, location, and severity, and recommending corrective strategies. Finally, once the devices are integrated with biological system, such as hiPSC-CM tissues in our demonstration, DeviceAgent autonomously analyzes the resulting electrophysiological recordings. This includes extracting cardiac extracellular field potentials, curating signals, analyzing waveform features, and mapping population-level dynamics to evaluate device performance.

### DeviceAgent enables autonomous design of flexible bioelectronics

To demonstrate the capabilities of DeviceAgent, we applied it to the design of flexible bioelectronic interfaces for cardiac tissue integration^29,30,31^, an application that demands careful coordination of mechanical compliance and electrical performance. In our demonstration, the design objectives included accommodating tissue growth through mechanical flexibility and stretchability, enabling seamless integration, and preserving high-resolution electrophysiological recording with cellular-scale electrodes. These objectives were translated into three sets of parameters: (1) geometry parameters, including device boundary, electrode number and placement, and routing endpoint directions; (2) mesh parameters, including electrode diameter, interconnect width, passivation margin, and mesh shape; and (3) input/output (I/O) pad parameters, including pad height, width, and pitch (**Extended Data Fig. 1**).

Manually coordinating these parameters is labor-intensive and error-prone, especially during iterative refinement. For example, modifying electrode distributions requires not only redesigning electrode placements but also re-routing interconnects, updating passivation structures, and rechecking for layer alignment. Similarly, minor updates to I/O pad configurations or mesh architecture could affect the entire device hierarchy, requiring updates across multiple layers.

DeviceAgent streamlines this process through an agent-driven workflow that spans from high-level design specification to fabrication-ready CAD files. Users begin by providing geometric parameters, such as the number of electrodes, device boundary dimensions, placement strategy, and routing endpoints, which DeviceAgent translates into electrode and endpoint layouts (**Fig 2a**). After establishing the basic geometry, the agent requests mesh parameters, including electrode diameter, interconnect width, passivation margin, and a preferred mesh shape (e.g., straight, sine, or round). The agent then autonomously and dynamically executes Python code to construct mesh layouts, route connections between electrodes and endpoints while enforcing spacing and alignment constraints, and shape mesh (e.g., sine waves) to endow stretchability (**Fig 2b**). The agent could also generate diverse design configurations with different electrode placements and mesh geometries, automatically producing optimized mesh and routing solutions and complete device layouts with I/O pads (**Extended Data Fig. 2**).

**Fig. 2:**
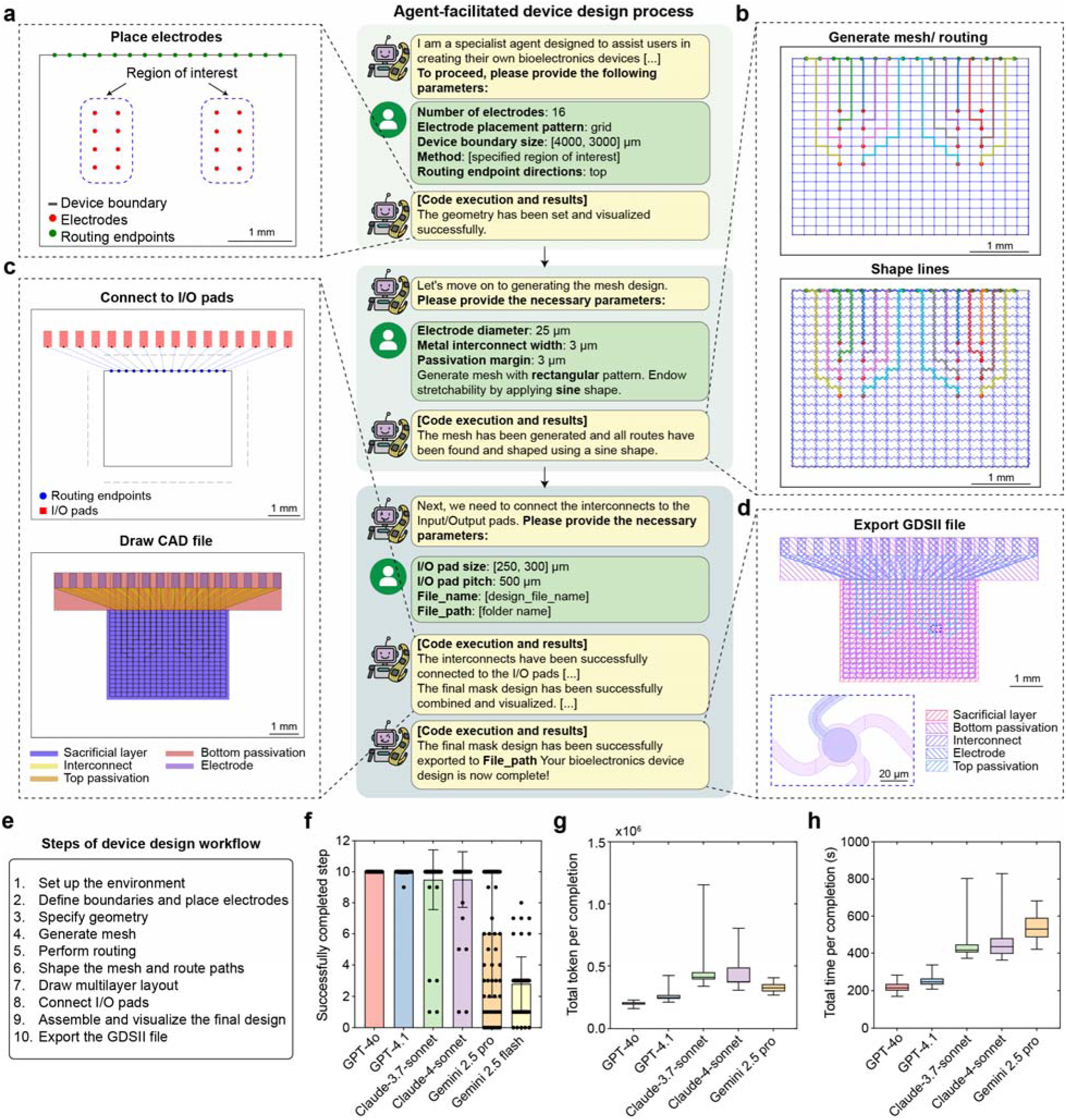
DeviceAgent-facilitated bioelectronics design. (**a-d**) Agent-facilitated device design workflow through a conversational interface: (a) Initial geometry setup based on user-specified parameters (number of electrodes, placement pattern, boundary size, routing directions). (b) Mesh generation with sine-shaped lines for stretchability. (c) I/O pad connection and CAD file generation showing the multilayer structure. (d) Final exported GDSII file with defined layers for sacrificial substrate, passivation, interconnects, and electrodes. (**e-h**) Benchmarking LLMs for automated layout generation: (e) 10-step design workflow used to benchmark each model’s ability to complete multistep layout generation tasks. (f) Number of successfully completed steps per trial for each model. Each dot represents one trial. (mean ± S.D., n=65-67). Trials that reached full completion (step 10) were selected for subsequent analyses. (g) Total token used per fully completed trials across models. (h) Total time per fully completed trial across models. Whiskers indicate minimum and maximum values. Sample sizes: GPT-4o (n = 67), GPT-4.1 (n = 66), Claude-3.7-Sonnet (n = 61), Claude-4-Sonnet (n = 60), Gemini 2.5 Pro (n = 29).

A notable strength of DeviceAgent is its ability to generate creative parametric mesh designs through few-shot in-context learning and structured code generation (**Extended Data Fig. 3**). With minimal prompting, it generates sophisticated mesh patterns such as wave-like distortions, which would be difficult and time-consuming to design manually. These parametric designs can be readily exported for finite element method (FEM) simulations to evaluate their mechanical behaviors. Once the mesh design is finalized, DeviceAgent automatically places I/O pads and generates multilayer device layouts for photolithography, including sacrificial, interconnect, electrode, and passivation layers with correct layer registration (**Fig 2c**). The final design is exported as a GDSII file for downstream use in photomask generation or direct-write lithography (**Fig. 2d**).

To evaluate the reliability of this multistep design pipeline, we benchmarked DeviceAgent across several LLMs, including Open AI GPT-4o, GPT-4.1, Anthropic Claude 3.7 and 4 Sonnet, and Google Gemini 2.5 Pro and Flash. The layout generation task was divided into 10 sequential steps, and each model was tested over multiple trials to determine how many steps it could complete per attempt (**Fig. 2e, f**). GPT-4o consistently completed all 10 steps, while GPT-4.1 completed them in all but one trial. Claude 3.7 and 4 Sonnet showed moderate success rates, followed by Gemini 2.5 Pro. Gemini 2.5 Flash failed to reach full completion in every attempt. We further compared performance metrics across models, including token usage and total time per completion (**Fig. 2g, h**) to show the trade-offs in efficiency and resource usage across LLMs. These performance differences likely reflect variations in reasoning robustness, tool-use reliability, and the ability to maintain context across extended workflows.

This modular, reproducible workflow accelerates the development of custom bioelectronic interfaces and reduces the time and expertise required for iterative design. By transforming the complex, manual process of layout generation into an agent-guided, conversational interaction, DeviceAgent lowers the technical barriers and improves accessibility, speed, and precision in bioelectronics design.

### DeviceAgent enables knowledge-driven nanofabrication protocol generation

Following device design, a critical next step in bioelectronics development is selecting compatible materials and developing a reliable layer-by-layer fabrication protocol. This process typically requires deep expertise in materials science and nanofabrication. For example, some materials are sensitive to high temperatures, while others require high temperature for curing. As a result, thermally sensitive layers must be patterned after those that require high-temperature processing. Chemical orthogonality also needs to be considered, since certain materials may swell or dissolve when exposed to solvents used in subsequent steps. These constrains necessitate careful sequencing of fabrication steps and often call for alternative patterning strategies. Even for experienced researchers, mastering this process requires years of practical experience and highly specialized expertise.

DeviceAgent addresses these challenges through autonomous protocol generation that integrates structured domain knowledge with real-time literature retrieval (**Fig. 3a**). The agent interprets user intent and retrieves relevant information on materials and fabrication procedures from both a curated local knowledge database and online literature sources (e.g., SerpAPI queries). It then synthesizes context from all available sources through a hybrid reasoning process and generates a detailed, step-by-step fabrication protocol using a dedicated guide-generation tool. Each protocol specifies materials, associated fabrication methods, and process constraints, ensuring layer compatibility and reproducibility.

**Fig. 3:**
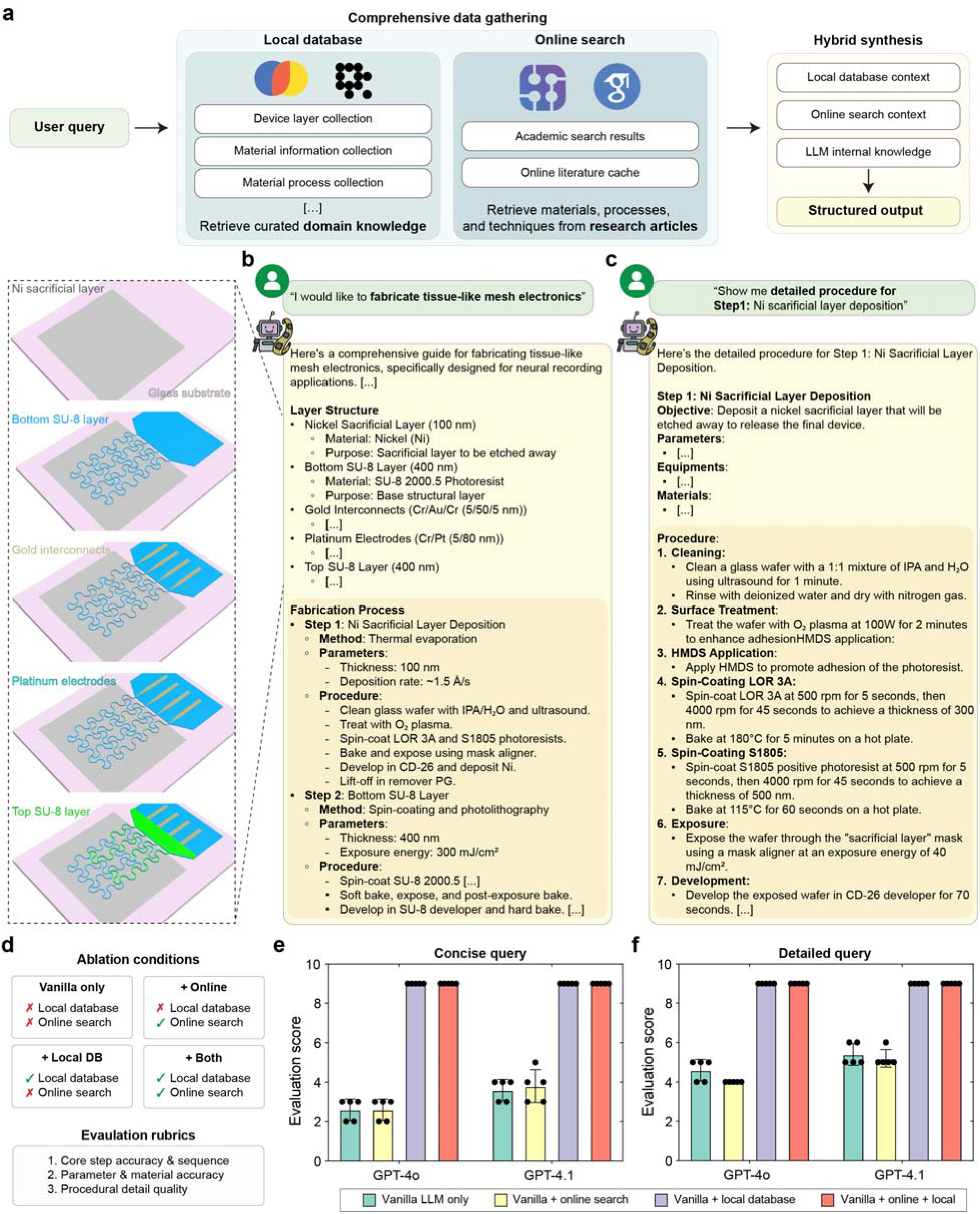
Knowledge-integrated nanofabrication protocol generation. (**a**) Schematic showing architecture of DeviceAgent’s protocol generation system, which integrates user queries with a curated local database and online literature sources to generate comprehensive fabrication instructions. (**b-c**) Examples of generated protocols: (b) Response to a general fabrication query, showing a complete layer structure and process flow. (c) Detailed procedure for a specific fabrication step with precise specifications. (**d**) Description of four ablation configurations used to evaluate protocol generation performance: Vanilla only (no external context), + Online (with online search context), + Local DB (with local database context), and +Both (with both online and local database context). Models generated protocols using either a simple or detailed version of the same query and evaluated using three rubrics. (**e-f**) Evaluation scores across different language models and query types: (e) simple query, and (f) detailed query. (mean ± S.D., n=5).

As an example, when prompted with “*I would like to fabricate tissue-like mesh electronics,*” DeviceAgent produced a detailed multilayer protocol (**Fig. 3b**). The output included material assignments–such as nickel (Ni) for the sacrificial layer, SU-8 for passivation, chromium/gold (Cr/Au) for interconnects, and platinum (Pt) for electrodes–followed by a step-by-step process detailing methods, key parameters, and procedural instructions. The agent also responds to more targeted queries. When asked “*Show me the detailed procedure for Ni sacrificial layer deposition,*” DeviceAgent generated a structured protocol that included objectives, required materials and equipment, and stepwise procedures spanning substrate cleaning, surface treatment, spin-coating, UV exposure, and development (**Fig. 3c**).

A key strength of DeviceAgent is its adaptability. It can adjust protocols to user requirements or equipment availability (**Extended Data Fig. 4**), for example by returning an electrochemical deposition protocol for PEDOT electrode modification (**Extended Data Fig. 4a**) or substituting polyimide for SU-8 as the substrate with updated spin coating and curing conditions (**Extended Data Fig. 4b**). DeviceAgent can also provide comparative material analysis. When asked to replace the Ni sacrificial layer, it evaluated alternatives such as aluminum, copper, and silicon dioxide based on their compatibility with downstream processes (**Extended Data Fig. 4c**).

To evaluate DeviceAgent’s architecture, we conducted ablation studies across four configurations: (1) LLM only (vanilla LLM), (2) LLM with online search, (3) LLM with local knowledge base, and (4) LLM with both local and online sources. Protocols were generated from both concise and detailed prompts and scored using rubric-based evaluation by an LLM judge (Methods; **Fig. 3d**). Access to the local knowledge base consistently improved performance, outperforming both vanilla LLMs and LLMs with online search only (**Fig. 3e**). While detailed prompts with materials and thickness information improved the baseline LLM output, they still fell short of protocols generated with structured knowledge integration (**Fig. 3f**). Notably, incorporating only online search module did not significantly enhance performance, as it currently relies on short snippets returned by SerpAPI, which lack the step-specific details available in full-text articles. This result suggests that precise multi-task protocol design requires packaging LLMs within a structured AI agent framework.

By combining structured internal knowledge, real-time literature retrieval, and model reasoning, DeviceAgent supports both standard and customized fabrication protocols. This flexibility makes it a powerful tool for developing and refining fabrication strategies in bioelectronics, particularly for multilayer systems where materials and process compatibility is critical.

### VLM-enabled visual inspection and troubleshooting in nanofabrication

Nanofabrication of bioelectronics is highly error-prone, where even minor defects can compromise device performance, reliability, and yield. In conventional workflows, researchers develop fabrication protocols, execute processes, and manually inspect devices to diagnose failures. While full automation is standard in semiconductor foundries for CMOS chip fabrication, such automation is difficult to implement in bioelectronics due to frequent design iterations, heterogeneous materials, and customized fabrication processes.

We demonstrated the capabilities of DeviceAgent in the fabrication of stretchable mesh electronics for cardiac organoid integration. These devices feature micrometer-scale patterns, high-density and narrow interconnects, and minimal passivation-to-interconnect margins, which is challenging for standard nanofabrication process. In such systems, seemingly minor issues such as trapped dust particles, micro-cracks, or residue contaminants can cause leakage, corrosion, or open circuits when exposed to biofluids. Layer-by-layer inspection is therefore essential for quality control, but manual review of microscopy images is labor-intensive and prone to overlooking subtle defects.

DeviceAgent addresses this challenge by integrating a VLM for automated visual inspection and defect reasoning (**Fig. 4a**, **b**). Using few-shot learning^32,33^ with annotated examples of both defect-free and defective bright-field (BF) microscopic images, the agent learns step-specific defect types and critical regions. BF images are processed through a vision encoder, paired with user-defined queries, and returned as structured outputs that include fabrication step identification, defect classification (e.g., dust, misalignment, scratches), spatial localization, severity scoring, and natural language explanations with recommended corrective actions.

**Fig. 4:**
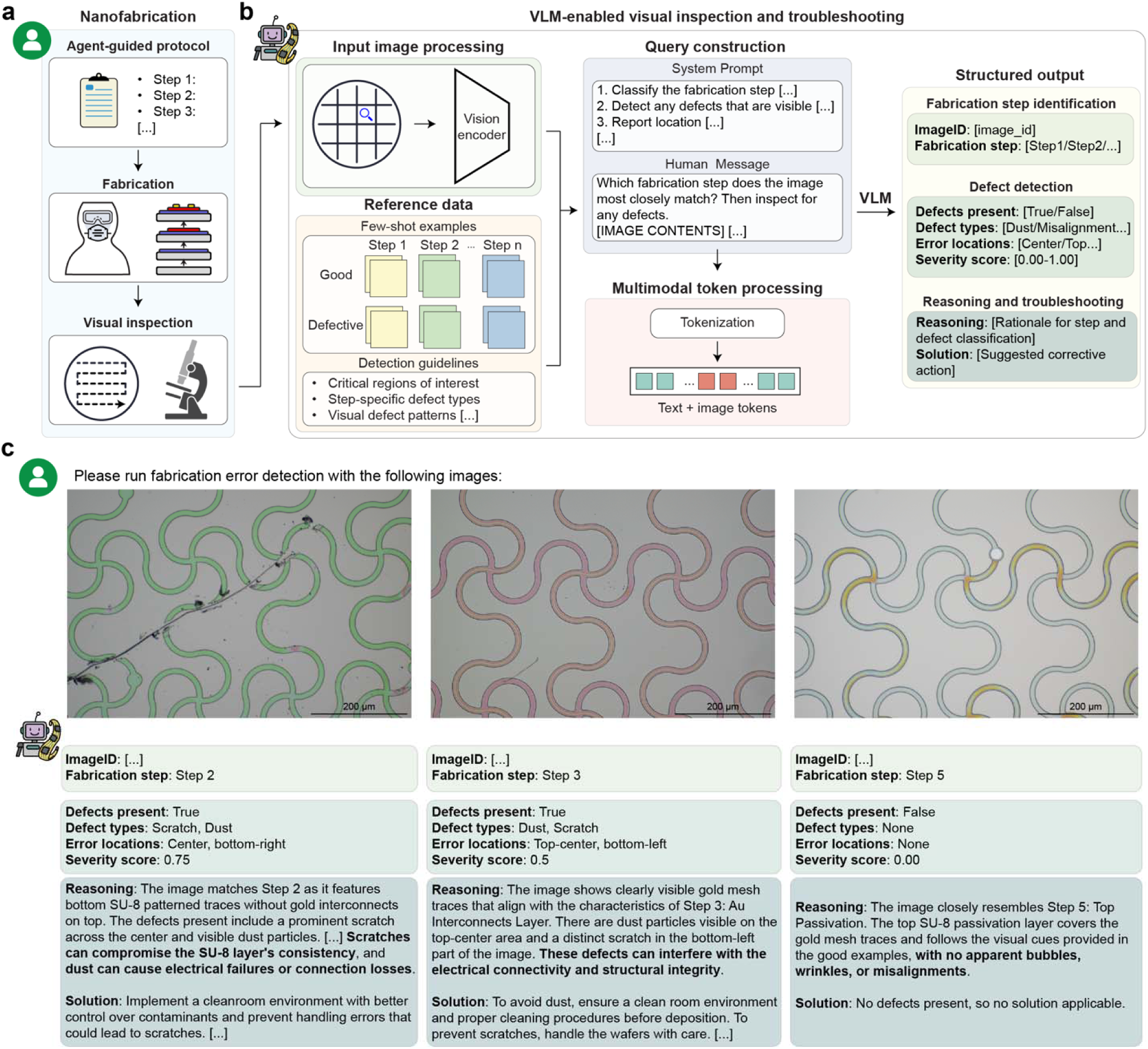
VLM-enabled visual inspection and troubleshooting in nanofabrication. (**a**) Schematic showing the nanofabrication workflow, including agent-guided protocol generation, fabrication processes, and visual inspection. (**b**) Schematic showing the architecture of the VLM-enabled visual inspection system: input image processing via a vision encoder, reference data with few-shot examples, query construction combining system prompts with human messages, and structured outputs including fabrication step identification, defect detection, reasoning, and suggested solutions. (**c**) DeviceAgent analyzing microscope images from three fabrication steps, demonstrating defect identification, severity assessment, reasoning about potential impacts, and recommended corrective actions.

We tested DeviceAgent on representative BF images from multiple fabrication steps of stretchable mesh nanoelectronics (**Fig. 4c**). For Step 2 (bottom SU-8 layer patterning), DeviceAgent accurately identified scratch and dust defects located in the center and bottom-right of the image, assigning high severity scores and reasoning that they could disrupt layer consistency and electrical connectivity. Recommended actions included stricter contamination control and improved handling protocols. For Step 3 (Cr/Au interconnects patterning), DeviceAgent detected both dust and scratches with moderate severity, warning of potential electrical connectivity and structural issues and suggesting enhanced cleaning and careful wafer handling. In contrast, an image from Step 5 (top SU-8 passivation) was correctly identified as defect-free with a severity score of 0.00, accompanied by reasoning consistent with reference examples.

Unlike conventional machine-learning approaches that require extensive, task-specific training^34^, this VLM-based framework leverages pretrained models and few-shot in-context learning, enabling rapid adaptation to new defect types, fabrication processes, and design variations with minimal user supervision. This makes DeviceAgent particularly well-suited for fast-paced prototyping and experimental environments, where flexible error detection and rapid feedback are essential.

### DeviceAgent evaluates bioelectronics performance

The final step in bioelectronics development is to evaluate device performance in biological experiments. In our demonstration, we integrated stretchable mesh electronics with hiPSC-derived cardiac tissues (**Fig. 5a-c**). Following release from the substrate via Ni sacrificial layer etching, devices were embedded within a Matrigel matrix and seeded with hiPSC-derived cardiomyocytes (**Extended Data Fig. 5**). These tissue-embedded electronics were then used to record electrophysiological activity from the tissues (**Fig. 5d, e**).

**Fig. 5:**
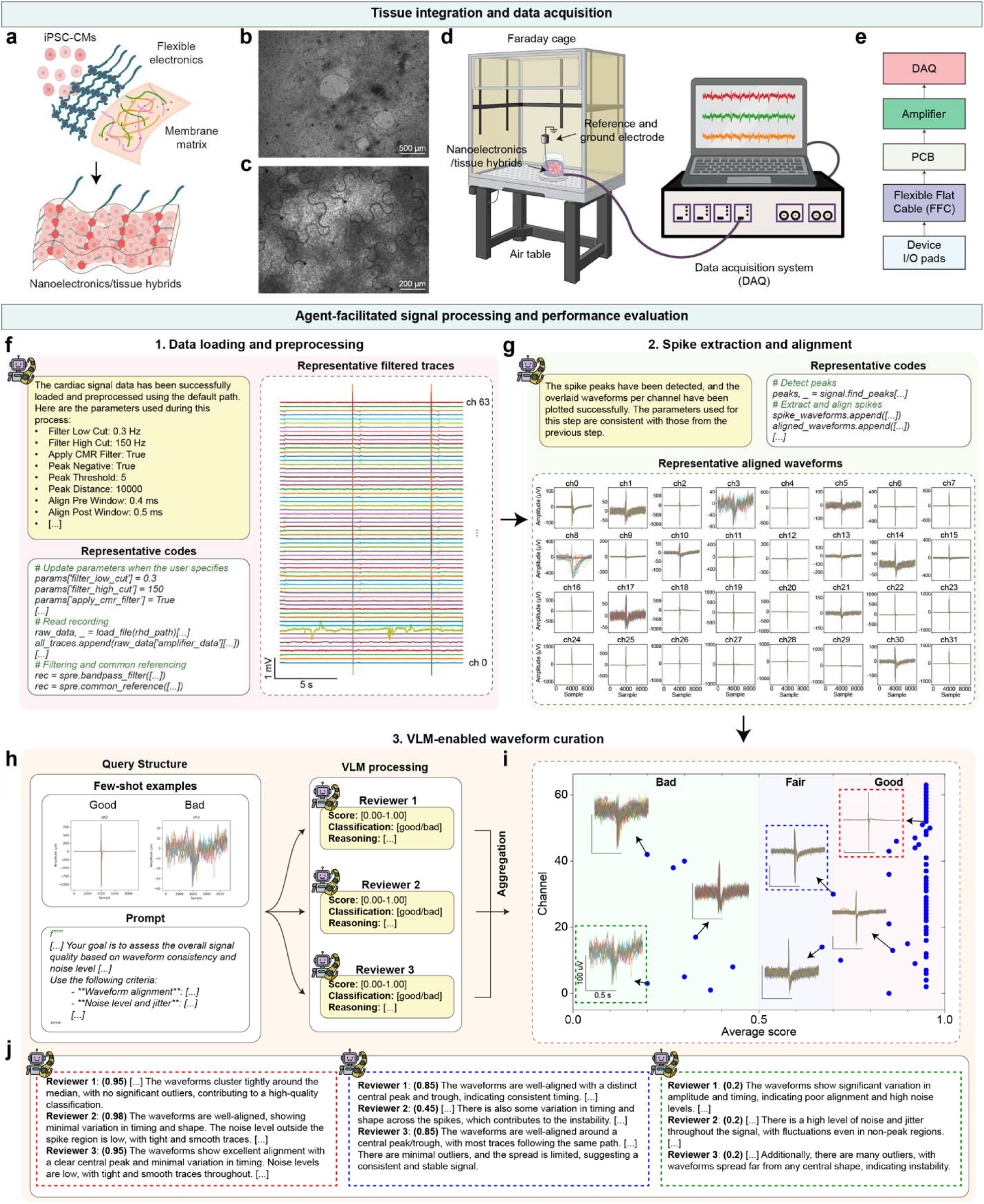
Tissue integration and agent-enabled performance evaluation. (**a-e**) Tissue integration and data acquisition: (a) Schematic of flexible electronics integrated with hiPSC-derived cardiomyocytes (hiPSC-CMs) using Matrigel to form tissue-electronics hybrid. (b-c) Microscopy images of (b) hiPSC-CMs and (c) the integrated tissue-electronics interface. (d) Schematic illustrating the experimental setup with a Faraday cage, air table, and data acquisition system. (e) Schematic showing hierarchical connection system from device I/O pads to data acquisition system. (**f-j**) Agent-facilitated signal processing workflow: (f) Data loading and preprocessing with agent-generated code and trace visualization. (g) Spike extraction and alignment with representative waveforms. (h) VLM-based waveform curation system using in-context few-shot examples, a multimodal prompt, and three independent AI reviewers. (i) Visualization of waveform quality assessments with channels categorized as good, fair, or bad. (j) Detailed reasoning provided by each AI reviewers for representative waveform classifications.

DeviceAgent autonomously validated the recording quality and the integrity of the tissue-device interface. Leveraging its code generation capability, the agent produced Python scripts to load raw recording files and apply standard preprocessing steps such as bandpass filters and common-mode referencing with either default or user-specified parameters (**Fig. 5f**). Preprocessed signals were then passed to a waveform extraction module that detected and aligned spikes across electrode channels (**Fig. 5g**).

A key innovation of using agentic framework is its automated waveform quality curation, a process that is traditionally manual and subjective^35^. Instead of relying on researchers to assess signal quality based on consistency, noise levels, and signal-to-noise ratios, DeviceAgent employs a VLM-based evaluation framework inspired by previous work in neural computation^15^. Using few-shot in-context learning and an ensemble learning approach^36^, the agent classifies waveforms directly from images. Few annotated examples categorized as “good” or “bad” are provided to prime the model (**Fig. 5h**). For each channel, three independent VLM-based “reviewers” assess waveform quality, assign scores, and generate textual explanations for their classifications. A final consensus label (e.g., good, fair, bad) is then derived by aggregating the assessments (**Fig. 5i**), with reasoning documented for transparency (**Fig. 5j**).

After identifying high-quality electrode channels, DeviceAgent performs downstream analyses to quantify spatiotemporal electrophysiological dynamics captured by the device. For example, it generated heatmaps of spike latency across channels during a single cardiac beat (**Extended Data Fig. 6a**), revealing spatial propagation patterns of electrical activity. It also extracted and visualized waveform features such as amplitude, maximum dV/dt, and half-width (**Extended Data Fig. 6b**), enabling quantitative comparisons across electrodes, experimental conditions, or time points.

By integrating tissue-electronics interfaces with automated signal processing, DeviceAgent provides a comprehensive, end-to-end framework from design to functional validation. Its ability to generate and execute code, curate signals via multi-agent review, and extract interpretable features significantly reduces the expertise barrier, accelerates analysis, and enhances reproducibility through standardized, well-documented workflows.

## Discussion

In this study, we introduced DeviceAgent, a multimodal AI agent that redefines the bioelectronics research workflow by integrating LLMs, VLMs, and domain-specific tools for device design, fabrication protocol generation, visual inspection, and signal analysis. By connecting previously isolated domains–materials science, electrical engineering, nanofabrication, and biology–DeviceAgent transforms an expertise-intensive process into a human-AI collaborative pipeline that lowers technical barriers and enhances accessibility. A key feature of DeviceAgent is its transparent and interpretable reasoning, which provides clear justifications for material selections, defect diagnoses, and signal quality assessments, thereby improving both user confidence and reproducibility.

We demonstrated DeviceAgent’s capabilities through the development of stretchable mesh electronics for cardiac organoid integration and measurement. Its conversational interface enables rapid, customized layout generation without manual CAD drafting. Protocol generation combines structured domain knowledge with real-time literature retrieval to produce fabrication workflows tailored to material and process constraints. VLM-powered visual inspection enables few-shot defect detection, adaptable to new defect types without large training datasets–an advantage for prototyping and iterative experimentation. For electrophysiological recordings, DeviceAgent automates preprocessing, signal curation, and feature extraction through a multi-reviewer VLM framework, delivering standardized, interpretable analyses. Together, these capabilities establish a comprehensive pipeline–from device design to functional validation–that accelerates development cycles and enhances reproducibility.

Compared with conventional frameworks, DeviceAgent offers several advantages. First, it lowers the expertise barrier by providing domain-specific guidance at each stage. Second, it accelerates iteration through continuous feedback and troubleshooting. Third, it maintains human oversight at critical decision points while automating repetitive or technically demanding tasks. Fourth, it leads to a more integrated workflow, where insights from later stages (e.g., fabrication issues or signal quality) can directly inform earlier stages (e.g., design modifications or protocol adjustments). Together, these features create a more accessible, efficient, and adaptive research process.

Although this study focused on establishing the technical framework of DeviceAgent, its broader scientific impact lies in advancing our understanding of bio-electronics interactions. Cyborg organoid models, in particular, offer a unique opportunity to systematically investigate how device geometry shapes tissue integration. By rapidly iterating mesh and interconnect designs and coupling them with automated evaluation of electrophysiological recordings, DeviceAgent can enable large-scale studies that reveal how the architectural feature of devices affects cellular organization, tissue maturation, and functional development. Such studies, which are impractical with manual design and characterization, could yield fundamental insights into the design principles that govern seamless integration between synthetic devices and living systems. Beyond geometry, integrating biological data with fabrication parameters may uncover correlations between material selections, device performance, and tissue responses–guiding the development of personalized or application-specific bioelectronic interfaces. In this way, DeviceAgent serves not only as an accelerator for device prototyping but also as a scientific instrument for uncovering the principles that underlie seamless biointegration and function, with implications for regenerative medicine and implantable systems.

At present, DeviceAgent is scoped to a specific application focused on serpentine mesh electronics for integration with hiPSC-derived tissues. This focus allowed us to direct our efforts toward building a human-AI-in-the-loop pipeline for an end-to-end demonstration. However, broader applications such as neural probes, wearable sensors, or implantable systems will require incorporation of application-specific design rules, fabrication processes, and data analysis methods. Future extensions could integrate finite element method (FEM) and reinforcement learning to autonomously optimize device geometries, or adopt modular strategies conceptually analogous to electronic design automation (EDA) in integrated circuit design, where standardized components and hierarchical abstractions support automated layout generation and validation^9^.

Several limitations remain. Protocol generation is reliable for familiar architectures but less robust for unconventional layer stacks or materials. Expanding capability will require broader knowledge integration, either through curated expert databases or full-text retrieval from scientific literature. Such workflow would involve multi-step processes for identifying relevant articles, parsing content, and extracting detailed experimental methods. While computationally demanding and potentially exceeding the input limits of current models, this approach could greatly improve the accuracy and specificity of generated protocols as model capacity and efficiency advance.

VLM-based defect inspection currently supports optical imaging but could be extended to multimodal characterization, including profilometry, electrical measurement, or optical spectroscopy. Signal analysis is optimized for cardiac electrophysiology but could be expanded to other modalities such as neural recordings, calcium imaging, or spatial transcriptomics through modular integration. Beyond fabrication and analysis, DeviceAgent may eventually support upstream processes such as protocol recommendation for cell culture and differentiation.

Together, these advances illustrate the potential for AI-augmented scientific workflows that remain grounded in domain expertise while increasing speed, reproducibility, and adaptability.

As platforms like DeviceAgent evolve, we anticipate that human-AI collaboration will become an increasingly central paradigm in bioelectronics research and broader scientific disciplines.

## Methods

### DeviceAgent backend framework

The backend infrastructure of DeviceAgent was developed using LangChain and LangGraph, which provide an orchestration layer for agent-based workflows driven by large language models (LLMs). LangChain manages prompt construction, tool invocation, and memory handling, while LangGraph enables structured control flow and state tracking across multi-step reasoning tasks. The system integrates multimodal LLMs, including GPT-4o and Claude 3.7 Sonnet, selected for their advanced reasoning, code generation, and vision–language capabilities. These models were used without fine-tuning to preserve their general-purpose adaptability across domains.

The platform supports memory-aware interactions, multimodal input processing, and tool-based execution for complex tasks such as device design, fabrication protocol generation, vision-language inspection, and electrophysiological signal analysis. Agent workflows follow a reasoning-action loop: (1) the agent receives a user prompt or contextual input, (2) selects and invokes appropriate tools, (3) generates and executes task-specific code, and (4) returns structured or visualized outputs. Tasks such as layout generation, image processing, and data analysis are handled through dynamically generated code, executed in real time using LangChain’s Python REPL tool. Outputs–including GDSII design layouts, image annotations, and waveform plots–are returned for further processing or user interaction.

### Design automation and layout generation

The device design agent implements an agentic workflow to automate the layout generation process for tissue-like mesh bioelectronics. Users define key design parameters such as electrode number, placement geometry, mesh configuration, and I/O routing endpoints. Based on these inputs, the agent generates and executes Python code to place electrodes, compute interconnect paths, apply stretchable geometries (e.g., sine-shaped traces), and assign I/O pad positions, while enforcing design constraints related to mechanical and electrical performance.

This process leverages prebuilt Python packages, including GDSPY and PHIDL, registered as tools within the LangChain framework. Executed via a Python REPL environment, these tools enable flexible and real-time layout generation. The agent supports both regular mesh designs and custom parametric patterns. Completed designs are automatically converted into predefined multilayer schematics and exported as GDSII files, ready for photolithography-based fabrication.

To benchmark the reliability of layout generation, we evaluated DeviceAgent across six LLMs: OpenAI GPT-4o, GPT-4.1, Anthropic Claude 3.7, Claude 4 Sonnet, and Google Gemini 2.5 Pro and Flash. Each model received the same system prompt, query, and task parameters to ensure consistent input conditions. The layout generation process was divided into 10 sequential steps, representing key phases of the workflow such as electrode placement, interconnect routing, and GDSII file generation. For each model, multiple trials were conducted, and the number of successfully completed steps, input/output token usage, and process time per trial were recorded to assess model efficiency.

### Fabrication protocol generation system

The fabrication protocol agent produces structured, step-by-step workflows by integrating information from both local and external sources. Internally, it queries a vector database (Chroma or LanceDB) containing curated nanofabrication protocols extracted from experimental records and publications. Each entry includes embedded metadata such as material stacks, processing parameters, and compatibility notes for semantic retrieval.

In parallel, the agent performs real-time literature searches via SerpAPI and Google Scholar, extracting fabrication strategies and materials data. Retrieved contexts from both sources are synthesized into a unified guide, organized by layer and formatted for direct use in micro/nanofabrication via a structured output tool.

To assess the contribution of each system component, we conducted ablation studies using a consistent user query across four configurations: (1) Vanilla LLM, relying solely on internal model knowledge; (2) Local RAG (Retrieval-Augmented Generation), combining the model with a curated local database; (3) Web RAG, integrating real-time Google Scholar search results; and (4) Hybrid RAG, incorporating both local and web-derived context. Each configuration was prompted with either a simple request (*“Generate a detailed, step-by-step fabrication protocol for tissue-like mesh electronics designed to integrate with a cyborg cardiac organoid.”*) or a detailed request (*“Generate a detailed, step-by-step fabrication protocol for tissue-like flexible and stretchable mesh electronics designed to integrate with a cyborg cardiac organoid. I plan to use a chamber attached on top of a glass wafer, incorporating a 5-layer mesh structure: a Ni sacrificial layer, SU-8 passivation layers, Au interconnects, and Pt electrodes, with an overall thickness of less than 2* μ*m. Please suggest any additional materials or processes required for the device.”*). Outputs were scored against a reference protocol^20^ using GPT-4o, evaluated according to rubrics assessing step accuracy and sequence, parameter and material accuracy, and procedural detail and fidelity. All agents operated under the same system prompt to ensure consistency.

### Vision-language-based fabrication error inspection and troubleshooting

To support quality control during fabrication, DeviceAgent incorporates a vision-language model (VLM) based on GPT-4o to analyze optical images from each fabrication step. Users provide a small number of annotated examples labeled as either defect-free or containing issues such as dust, scratches, cracks, or misalignment. Optional textual guidelines can be supplied to emphasize regions critical to device function. Images are processed via a vision encoder and combined with text prompts to form a multimodal input for the LLM. The agent returns diagnostic outputs including the defect type, location, severity score, and recommended mitigation strategies via a structured output tool.

### Signal processing module

The signal analysis agent processes extracellular electrical recordings from bioelectronic devices, with a focus on signals acquired from hiPSC-derived cardiomyocytes. Raw data are loaded and preprocessed using a configurable bandpass filter (default: 0.3-150 Hz). Spikes are extracted from each channel and aligned to the point of maximum dV/dt in the waveform.

To evaluate signal quality, the agent uses a VLM based on GPT-4o, trained via few-shot learning on waveform examples labeled as “good” or “bad.” For each channel, three independent VLM reviewers assign classification scores and provide justifications. These are aggregated into a consensus label (e.g., good, fair, bad) with accompanying rationale. This approach enhances reproducibility and reduces bias in signal selection.

High-quality channels are passed to downstream analyses, including spike latency mapping, which calculates timing differences across electrodes during individual cardiac beats, and waveform feature calculation, including amplitude, maximum dV/dt, and half-width.

### Frontend and user interaction

The user interface of DeviceAgent was developed using Streamlit^37^, providing an accessible and interactive web interface that supports both graphical input forms and conversational prompts. Users can input design parameters, microscopy images, or electrophysiological recording data files and receive multimodal outputs in real time.

A module selection panel allows users to choose between different agent functionalities–device design, fabrication protocol generation, vision-language inspection, and signal analysis–tailoring the agent to specific stages of the bioelectronics research workflow.

The frontend returns context-aware, multimodal outputs, including layout previews, fabrication protocols, curated waveform plots, and structured defect assessments. Visualizations are generated dynamically through agent-produced Python code, allowing users to explore results with minimal technical overhead. Session persistence is managed through Streamlit’s session state, maintaining continuity across multi-step design and analysis workflows. The frontend operates within the same runtime as the LangChain-LangGraph backend, enabling direct integration of agent logic and tool execution.

### Device fabrication

A soda lime glass wafer (University Wafer) was cleaned in a 1:1 IPA/H O solution using an ultrasonic bath for 1 min, rinsed with deionized water, and dried with nitrogen. The surface was treated with O plasma (100 W, 2 min), followed by sequential spin-coating of hexamethyldisilazane (HMDS, MicroChem), LOR 3A (MicroChem; 500 rpm for 5 s, then 4000 rpm for 45 s), and S1805 positive photoresist (MicroChem; 500 rpm for 5 s, then 4000 rpm for 45 s). The wafer was baked at 180 °C for 5 min (LOR 3A) and 115 °C for 60 s (S1805), UV-exposed at 40 mJ/cm², developed in CD-26 developer (Microposit) for 70 s, rinsed with IPA, and dried with N. A brief O plasma treatment (50 W, 30 s) was applied prior to thermal evaporation of a 100 nm Ni sacrificial layer (∼1.5 Å/s). Liftoff was achieved by overnight immersion in Remover PG (MicroChem), followed by IPA rinse and N drying. For the bottom SU-8 layer, the wafer was again plasma-treated (100 W, 2 min), spin-coated with SU-8 2000.5 (MicroChem; 500 rpm for 5 s, then 4000 rpm for 45 s), soft-baked (65 °C for 2 min and 95 °C for 4 min), UV-exposed at 300 mJ/cm², post-exposure baked (65 °C for 2 min and 95 °C for 2 min), and developed in SU-8 developer (MicroChem) for 60 s. After rinsing with IPA and drying, a hard bake was performed by ramping the temperature from 65 °C to 180 °C over 40 min and cooling slowly. For the Au interconnects, the wafer was treated with O plasma (50 W, 30 s), spin-coated with HMDS, LOR 3A, and S1805 as above, UV-exposed at 40 mJ/cm², and developed in CD-26. After another O plasma step (50 W, 30 s), a Cr/Au/Cr stack (5/50/5 nm) was deposited via e-beam evaporation (∼2.0 Å/s), and liftoff was carried out in Remover PG.

The Pt electrodes were patterned using the same process, followed by e-beam evaporation of Cr (5 nm, ∼2.0 Å/s) and Pt (80 nm, ∼1.5 Å/s), and subsequent liftoff. The top SU-8 passivation layer was then formed by spin-coating SU-8 2000.5, soft-baking (65 °C for 2 min and 95 °C for 4 min), UV exposure at 300 mJ/cm², post-exposure baking (65 °C for 2 min and 95 °C for 2 min), and development in SU-8 developer for 70 s. A final hard bake was performed by ramping from 65 °C to 190 °C over 40 min. Devices were diced using a dicing saw, and flexible flat cables (Molex) were soldered to the I/O pads using a flip-chip bonder at 180 °C for 30 s. A custom chamber was fabricated by cutting a 15 mL centrifuge tube (VWR) and sealing it to the wafer using Kwik-Sil silicone (World Precision Instruments), cured overnight at room temperature. Finally, the mesh device was released by performing an O plasma treatment (50 W, 30 s), adding Ni etchant (Transene) into the chamber for 2–4 hours, and thoroughly rinsing with deionized water.

### Cell culture and integration

#### Stem cell culture and cardiomyocyte differentiation

Human induced pluripotent stem cells (hiPSCs, IMR-90 line; WiCell Research Institute, Madison, WI, USA) were cultured and differentiated following previously established protocols. Cells were maintained on Matrigel-coated 6-well plates in Essential 8 medium (Gibco), with daily medium changes. Passaging was performed every four days using 5 mM EDTA (Invitrogen). Cardiomyocyte differentiation was initiated at ∼80% confluency using RPMI 1640 medium supplemented with 2% B27 without insulin (Gibco) and 12 µM CHIR99021 (BioVision) for 24 hours. This was followed by 48 hours in the same medium without CHIR99021, then treatment with 5 µM IWR-1 (Cayman Chemical) for an additional 48 hours. Cells were then maintained in RPMI 1640 with 2% B27 without insulin for recovery and subsequently cultured in RPMI 1640 with 2% B27. Spontaneous contractions were typically observed 2–3 days after B27 supplementation. Medium was replaced every 48 hours.

#### Device integration with hiPSC-derived cardiomyocytes

Devices were prepared for cell seeding by rinsing twice with 1× PBS, then incubated overnight in 50 µg/ml Poly-D-lysine hydrobromide (Sigma-Aldrich), rinsed with deionized water, and coated with Matrigel (100 µg/mL) at 37 °C for 1 hour. After chilling, 50 µL of 10 mg/mL Matrigel was applied under the device to create a tissue supporting layer, followed by 30 min of solidifying at 37 °C. hiPSC-derived cardiomyocytes were dissociated using 0.05% Trypsin-EDTA (Gibco), and viability assessed via trypan blue staining and TC20 Automated Cell Counter (BioRad). Approximately 3–4 million cells were seeded onto the Matrigel-coated devices. ROCK inhibitor (10 µM) was included during the first 24 hours post-integration and omitted thereafter. Culture medium was replaced every 48 hours.

### Electrophysiological recording

Electrophysiological recordings were acquired using the Intan RHD system. Device I/O pads were connected to Intan RHD headstages via a customized printed circuit board (PCB), with a platinum electrode immersed in the culture medium serving as both ground and reference. Recordings were conducted for 5 minutes inside a custom Faraday cage at a sampling rate of 10 kHz.

## Code and data availability

Code and data will be made publicly available on Github upon publication.

## Acknowledgements

J. Liu acknowledges the support from the National Institutes of Health (NIDDK 1DP1DK130673, NLM 5R01LM014465, and NHLBI 1R33HL175683-01). J. Lee acknowledges the support from Kwanjeong Educational Foundation.

## Author contributions

J. Lee, Z. L., W. W., and J. Liu conceived the idea. J. Lee, Z. L., and J. B. developed the DeviceAgent methodology. J. Lee and A. J. L. fabricated the devices. W. W. differentiated the cells and integrated the tissues with the devices. J. Lee and W. W. performed the electrophysiological recordings. J. Lee and W. W. prepared the figures. J. Lee, Z. L., and W. W. wrote the initial draft of the manuscript. All authors contributed to critical discussions, data interpretation, and figure refinement. J. Liu supervised the study.

---

**Extended Data Fig. 1:**
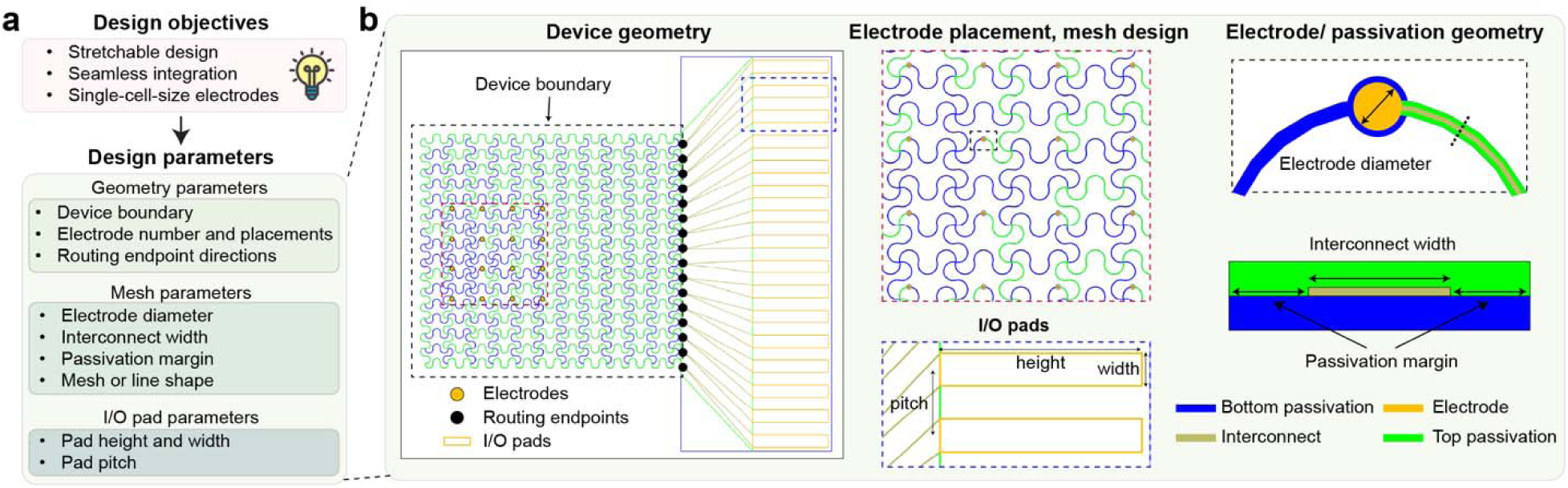
Design parameters of tissue-like mesh electronics. (**a**) Design objectives–stretchable architecture, seamless tissue integration, and single-cell scale electrodes– are translated into specific parameters across three categories: geometry parameters (device boundary, electrode placement, and routing endpoints); mesh parameters (electrode diameter, interconnect width, passivation margin, and meshline shape); and I/O pad parameters (height, width, and pitch). (**b**) Detailed components of bioelectronics design showing device geometry with electrode and routing endpoint placement, mesh design with interconnect patterns, and electrode/passivation geometry with critical dimensions.

**Extended Data Fig. 2:**
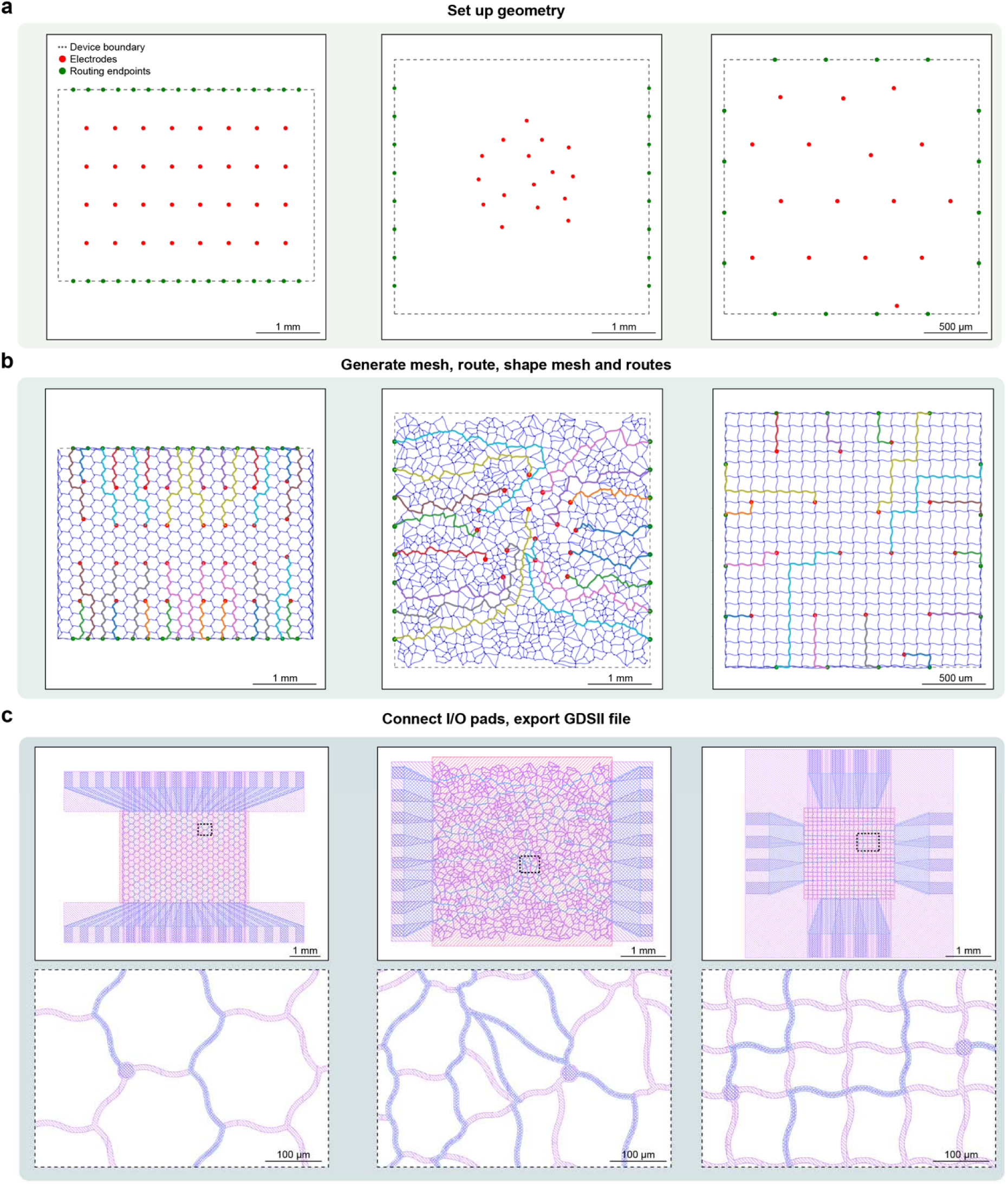
DeviceAgent-enabled design capabilities for diverse electrode arrangements. (**a**) Initial geometry setup for three different electrode arrangements: a regular grid pattern (left), a random distribution (middle), and a sparse arrangement (right). (**b**) Mesh and routing solutions generated by DeviceAgent for each electrode configuration, showing adaptation to different spatial distributions. (**c**) Complete device layouts with I/O pads for all three designs, with zoomed-in views of the mesh structures highlighting detailed interconnect patterns.

**Extended Data Fig. 3:**
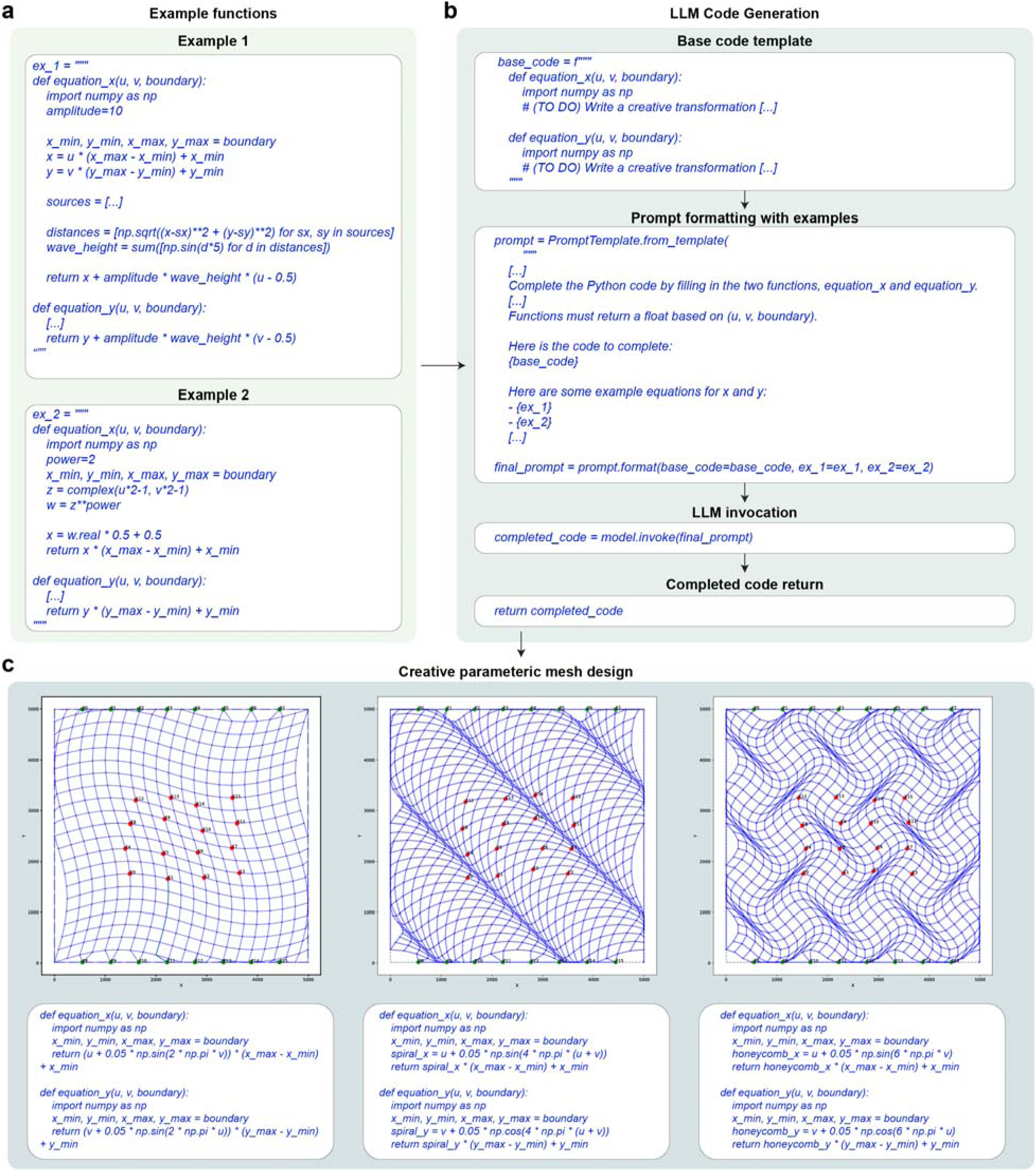
DeviceAgent-enabled code generation for creative parametric mesh designs. (**a**) Example functions used for few-shot learning, showing wave-like patterns (Example 1) and complex geometric transformations (Example 2). (**b**) Workflow of LLM-based code generation, starting with a base template, formatting a prompt with examples, invoking the LLM, and returning the completed code. (**c**) Creative parametric mesh designs generated by DeviceAgent, each corresponding to its respective mathematical transformation functions.

**Extended Data Fig. 4:**
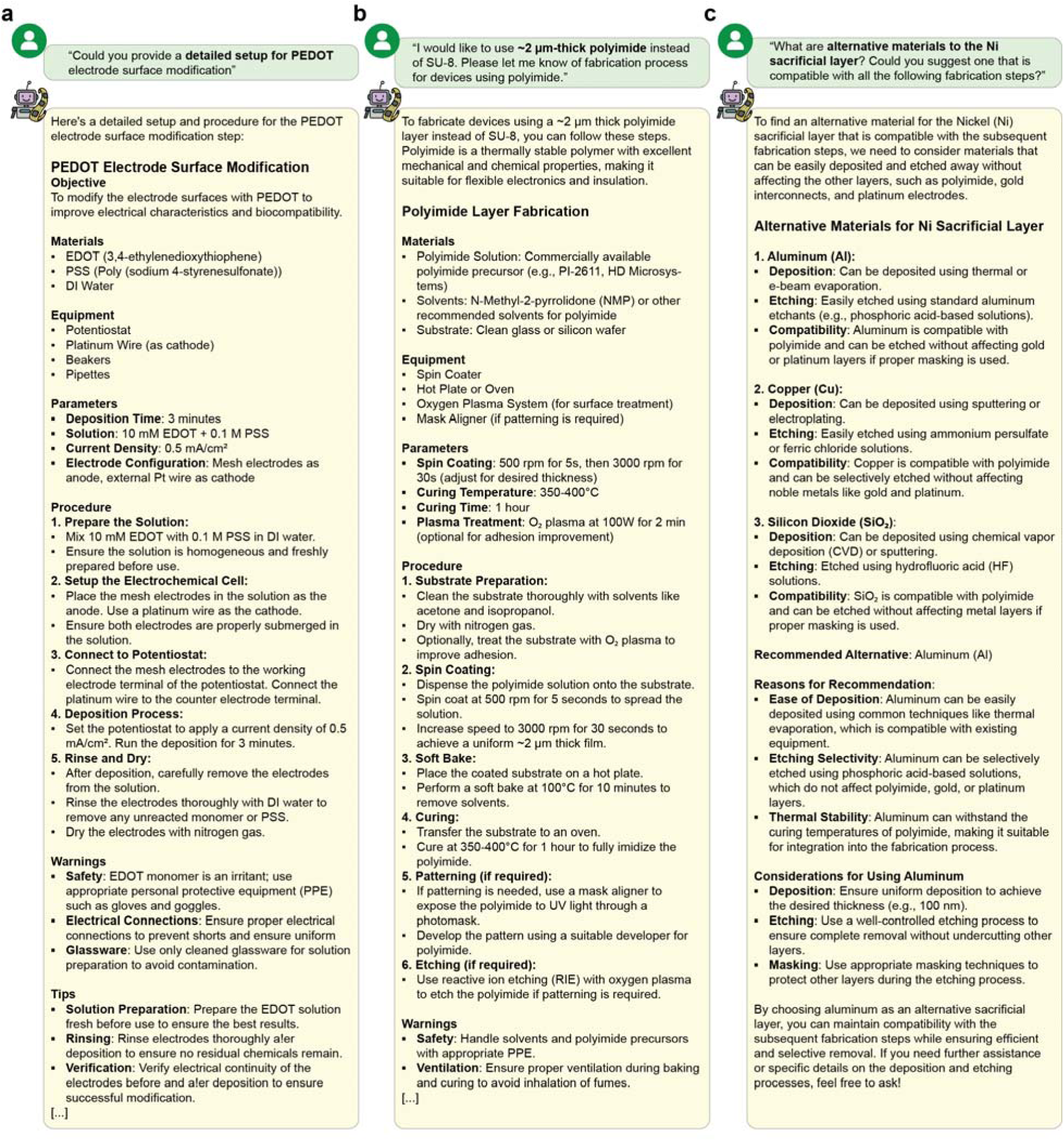
DeviceAgent-enabled adaptive protocol generation. (**a**) Detailed protocol for PEDOT electrode surface modification generated in response to a user query. The protocol includes comprehensive information on the objective, materials, equipment, parameters, and step-by-step procedures for electrochemical deposition. (**b**) Alternative fabrication protocol for using polyimide instead of SU-8, demonstrating DeviceAgent’s ability to adapt to material substitutions. The protocol specifies spin-coating parameters, curing conditions, and processing steps tailored to the properties of polyimide. (**c**) Comparative analysis of alternative materials to replace nickel sacrificial layer, with DeviceAgent evaluating aluminum, copper, and silicon dioxide based on deposition methods, etching processes, and compatibility with other device layers. The agent provides a reasoned recommendation for aluminum, including specific considerations for implementation.

**Extended Data Fig. 5:**
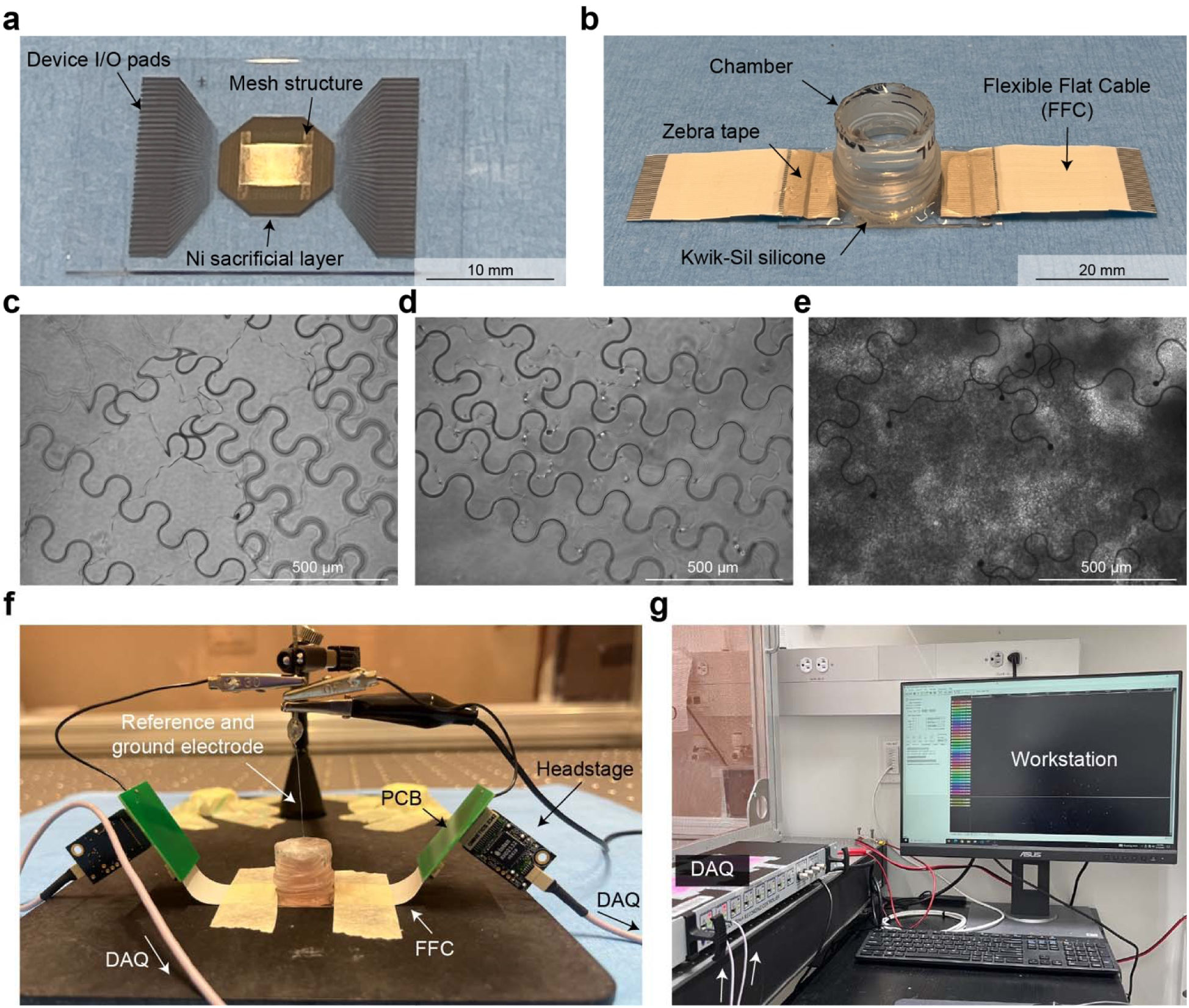
Device post-processing, tissue integration, and electrophysiology recording setup. (**a**) Representative image of a single mesh nanoelectronics device after dicing, showing the I/O pads, central mesh structure, and underlying Ni sacrificial layer. (**b**) Representative assembled device ready for cell culture, connected to a flexible flat cable (FFC) via zebra tape and enclosed within a silicone-sealed chamber. (**c-e**) Sequential steps of tissue integration: (c) Device after release via Ni sacrificial layer etching. (d) Matrigel membrane matrix applied beneath the device. (e) hiPSC-CMs seeded onto the device/Matrigel layer. (**f**) Electrophysiology recording setup showing connections from the device to the data acquisition (DAQ) system via a customized PCB and headstage. A platinum (Pt) electrode serves as both reference and ground within the recording chamber. (**g**) Workstation with DAQ system for real-time data acquisition and signal visualization.

**Extended Data Fig. 6:**
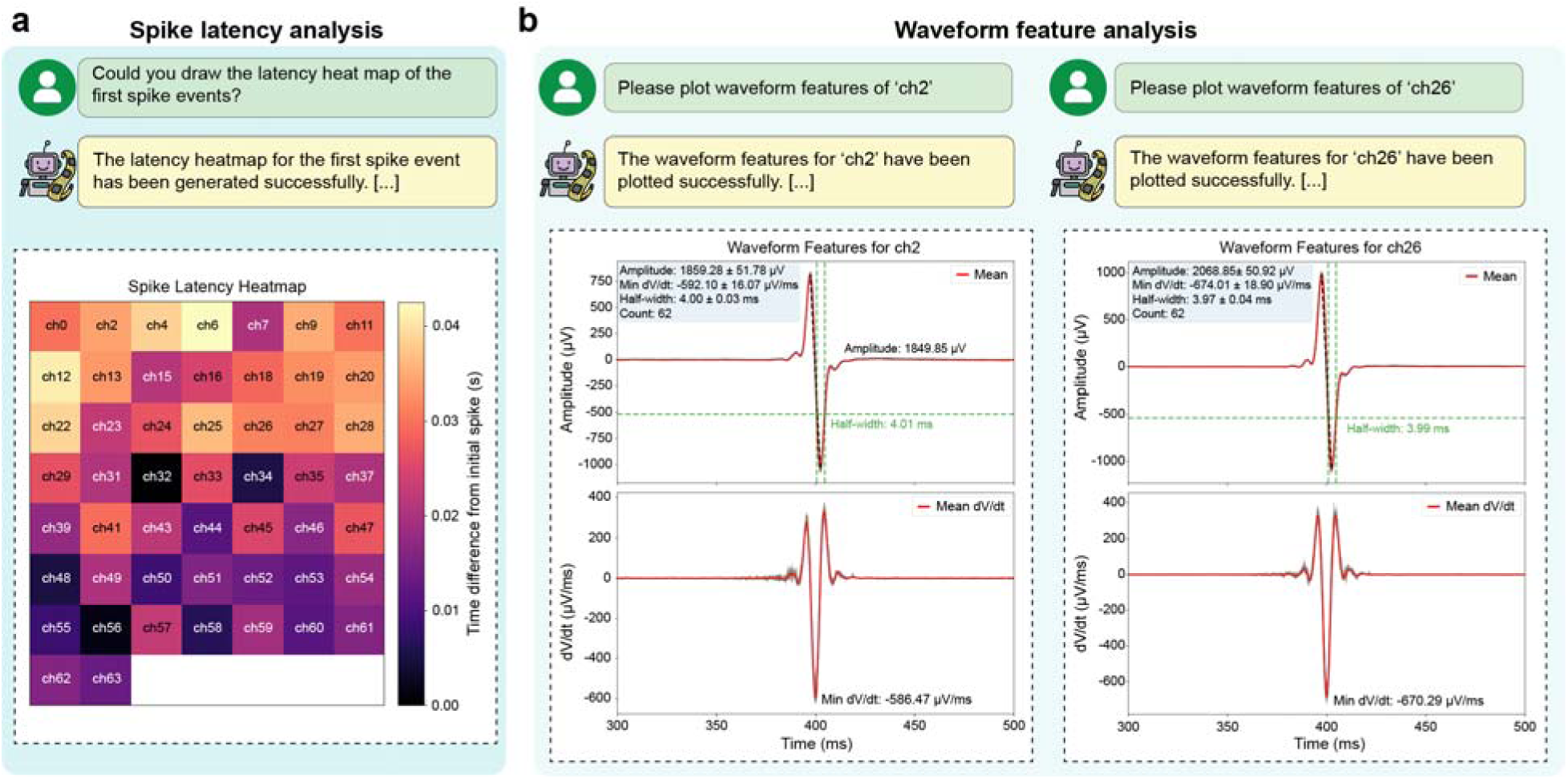
Electrophysiological analysis capabilities of DeviceAgent. (**a**) Spike latency analysis showing a heat map visualization of temporal delays from the initial spike across multiple recording channels, revealing the propagation pattern of electrical activity across the electrode array. (**b**) Waveform feature analysis for representative channels, displaying amplitude, maximum dV/dt, half-width, and spike count metrics with corresponding visualizations, enabling quantitative comparison across recording sites.

## References

1. Zhao, C. et al. Skin-inspired soft bioelectronic materials, devices and systems. Nat. Rev. Bioeng. 2, 671–690 (2024).

2. Song, E. et al. Materials for flexible bioelectronic systems as chronic neural interfaces. Nat. Mater. 19, 590–603 (2020).

3. Kim, H. J. et al. Materials design and integration strategies for soft bioelectronics in digital healthcare. Nat. Rev. Mater. 10, 654–673 (2025).

4. Zhao, S. et al. Tracking neural activity from the same cells during the entire adult life of mice. Nat. Neurosci. 26, 696–710 (2023).

5. Sheng, H. et al. Brain implantation of soft bioelectronics via embryonic development. Nature 642, 954–964 (2025).

6. Zhang, Y. et al. Millimetre-scale bioresorbable optoelectronic systems for electrotherapy. Nature 640, 77–86 (2025).

7. Trouillet, A. et al. High-resolution prosthetic hearing with a soft auditory brainstem implant in macaques. *Nat*. Biomed. Eng. 9, 1403–1417 (2025).

8. Milekovic, T. et al. A spinal cord neuroprosthesis for locomotor deficits due to Parkinson’s disease. Nat. Med. 29, 2854–2865 (2023).

9. Huang, G. et al. Machine Learning for Electronic Design Automation: A Survey. ACM Trans. Des. Autom. Electron. Syst. 26, 1–46 (2021).

10. Minaee, S., et al. Large Language Models: A Survey. arXiv preprint arXiv:2402.06196 (2024).

11. Liu, H., et al. Visual Instruction Tuning. arXiv preprint arXiv:2304.08485 (2023).

12. Bubeck, S., et al. Sparks of Artificial General Intelligence: Early experiments with GPT-4. arXiv preprint arXiv:2303.12712 (2023).

13. Kojima, T., et al. Large Language Models are Zero-Shot Reasoners. arXiv preprint arXiv:2205.11916 (2022).

14. Yao, S., et al. ReAct: Synergizing Reasoning and Acting in Language Models. arXiv preprint arXiv:2210.03629 (2022).

15. Lin, Z. et al. Spike sorting AI agent. bioRxiv preprint 10.1101/2025.02.11.637754 (2025).

16. Lin, Z. et al. Spatial transcriptomics AI agent charts hPSC-pancreas maturation in vivo. bioRxiv preprint 10.1101/2025.04.01.646731 (2025).

17. Aljović, A. et al. An autonomous AI agent for universal behavior analysis. bioRxiv preprint 10.1101/2025.05.15.653585 (2025).

18. Boiko, D. A. et al. Autonomous chemical research with large language models. Nature 624, 570–578 (2023).

19. Li, Q. et al. Cyborg Organoids: Implantation of Nanoelectronics via Organogenesis for Tissue-Wide Electrophysiology. Nano Lett. 19, 5781–5790 (2019).

20. Lin, Z. et al. Cyborg organoids integrated with stretchable nanoelectronics can be functionally mapped during development. Nat. Protoc. 20, 2528–2559 (2025).

21. OpenAI, et al. GPT-4o System Card. Available at: https://cdn.openai.com/papers/GPT-4o-System-Card.pdf (2024).

22. Anthropic. Claude 3.7 Sonnet. Available at: https://www.anthropic.com/news/claude-3-7-sonnet (2025).

23. Chase, H. LangChain. Available at: https://github.com/hwchase17/langchain (2023)

24. Wang, X. et al. Executable Code Actions Elicit Better LLM Agents. arXiv preprint arXiv:2402.01030 (2024).

25. Lewis, P., et al. Retrieval-Augmented Generation for Knowledge-Intensive NLP Tasks. arXiv preprint arXiv:2005.11401 (2020).

26. SerpAPI. SerpAPI. Available at: https://github.com/serpapi (2023).

27. Heitzmann, L. Gdspy: Python module for creating GDSII stream files. Available at: https://github.com/heitzmann/gdspy (2019).

28. McCaughan, A. N. et al. PHIDL: Python-based layout and geometry creation for nanolithography. J. Vac. Sci. Technol. B 39, 062601 (2021).

29. Lee, J. et al. In Situ Graphene-Seq: Spatial Transcriptomics and Chronic Electrophysiological Characterization of Tissue Microenvironments. bioRxiv preprint 10.1101/2025.03.25.645278 (2025).

30. Lin, Z. et al. Tissue-embedded stretchable nanoelectronics reveal endothelial cell– mediated electrical maturation of human 3D cardiac microtissues. Sci. Adv. 9, eade6460 (2023).

31. Li, Q. et al. Multimodal charting of molecular and functional cell states via in situ electro-sequencing. Cell 186 2002–2017 (2023).

32. Lampinen, A. K. et al. On the generalization of language models from in-context learning and finetuning: a controlled study. arXiv preprint arXiv:2505.00661 (2025).

33. Wei, J., et al. Chain-of-Thought Prompting Elicits Reasoning in Large Language Models. arXiv preprint arXiv:2201.11903 (2022).

34. Ren, Z. et al. State of the Art in Defect Detection Based on Machine Vision. Int. J. of Precis. Eng. Manuf.-Green Technol. 9, 661–691 (2022).

35. Wood, F. et al. On the variability of manual spike sorting. IEEE Trans. Biomed. Eng. 51, 912–918 (2004).

36. Wang, X. et al. Self-Consistency Improves Chain of Thought Reasoning in Language Models. arXiv preprint arXiv:2203.11171 (2022).

37. Streamlit. Streamlit. Available at: https://github.com/streamlit/streamlit (2023).

